# Reactivation-induced memory integration prevents proactive interference in perceptual learning

**DOI:** 10.1101/2022.09.01.506161

**Authors:** Zhibang Huang, Zhimei Niu, Sheng Li

**Affiliations:** School of Psychological and Cognitive Sciences, Peking University; Beijing Key Laboratory of Behavior and Mental Health, Peking University; PKU-IDG/McGovern Institute for Brain Research, Peking University; Key Laboratory of Machine Perception (Ministry of Education), Peking University; Department of Psychology, University of Texas at Austin

**Author notes:** Correspondence concerning this article should be addressed to Sheng Li, School of Psychological and Cognitive Sciences, Peking University, 5 Yiheyuan Road, Haidian, Beijing, 10087, China.

**Keywords:** long-term memory, reconsolidation, attentional suppression, perceptual training

## Abstract

We acquire perceptual skills through experience to adapt ourself to the changing environment. Accomplishing an effective skill acquisition is a main purpose of perceptual learning research. Given the often observed learning effect specificity, multiple perceptual learnings with shared parameters could serve to improve the generalization of the learning effect. However, the interference between the overlapping memory traces of different learnings may impede this effort. Here, we trained human participants on an orientation discrimination task. We observed a proactive interference effect that the first training blocked the second training at its untrained location. This was a more pronounced effect than the well-known location specificity in perceptual learning. We introduced a short reactivation of the first training before the second training and successfully eliminated the proactive interference when the second training was inside the reconsolidation time window of the reactivated first training. Interestingly, we found that practicing an irrelevant task at the location of the second training immediately after the reactivation of the first training could also restore the effect of the second training but in a smaller magnitude, even if the second training was conducted outside of the reconsolidation window. We proposed a two-level mechanism of reactivation-induced memory integration to account for these results. The reactivation-based procedure could integrate either the previously trained and untrained locations or the two trainings at these locations, depending on the activated representations during the reconsolidation process. The findings provide us with new insight into the roles of long-term memory mechanisms in perceptual learning.

## Introduction

Early theories of visual perceptual learning considered the improvement of visual functions through experience as a result of perceptual plasticity (Hochstein & Ahissar, 2002; Sagi & Tanne, 1994). A hallmark of perceptual learning that supported this proposal was the initially observed location, feature, or eye specificity of the learning effects (Ahissar & Hochstein, 1997; Ball & Sekuler, 1987; Crist et al., 1997; Fahle, 1997; Karni & Sagi, 1991; Shiu & Pashler, 1992). However, the accumulating evidence revealed by succeeding investigations pointed out that the underlying mechanism of perceptual learning is not purely perceptual. These studies have found that the learning specificity could also be explained by higher level factors such as restricted attentional processing (Donovan & Carrasco, 2018; Donovan et al., 2015; Xiao et al., 2008; Xiong et al., 2016) and modified reading-out of sensory information from decision areas (Dosher & Lu, 1998; Jia et al., 2018; Jia et al., 2020; Law & Gold, 2008). Not beyond our expectation, the roles of mnemonic processing in perceptual learning are drawing increasing attention as well. Recent investigations have suggested that perceptual learning effect is associated with the changed representation of visual stimuli in short-term memory (Jia et al., 2021) and refined consolidation process of long-term memory (Bang et al., 2018; Tamaki et al., 2020; Yang et al., 2022).

One of the ultimate goals of perceptual learning research is to promote its application in real-world (Deveau et al., 2014; Johnston et al., 2020; Sha et al., 2020). That would require the evidence of performance improvement in the contexts that are not identical to that of the training. Intuitively, this could be achieved by facilitating the generalization of the learning effect (Harris et al., 2012; Kattner et al., 2017). However, the widely observed learning specificity in perceptual learning was an obvious barrier against this idea. To address this issue, several procedures based on double training (Wang et al., 2014; Xiao et al., 2008), attentional cueing (Donovan & Carrasco, 2018; Donovan et al., 2015), or concept-related categories (Tan et al., 2019; Wang et al., 2016) have been proposed to improve the transfer of learning effect to untrained stimuli or retinal locations. These approaches, though successful in demonstrating the generalized learning effect, required other manipulations in addition to the trained task.

To extend the acquired learning effect to the untrained conditions, we could also conduct the training with modified parameters. The hope is that by sharing a subset of the parameters with the original training, the new training may require reduced effort to accomplish. However, interference is a ubiquitous phenomenon when multiple learnings with shared parameters take place (Anderson, 2003; Herszage & Censor, 2018; Postman & Underwood, 1973) and perceptual learning was not an exception (Bang et al., 2018; Been et al., 2011; Huang et al., 2022; Seitzt et al., 2005; Shibata et al., 2017; Yotsumoto et al., 2009). Particularly, several studies have shown that the interference can be observed between two trainings that were separated by at least one day, indicating that long-term memory mechanism was involved in and may provide the solution for the interference between multiple perceptual trainings (Bang et al., 2018; Huang et al., 2022).

It has long been suggested that the consolidation of learning-related memory traces is not a static process. Reactivating a consolidated memory can transfer it to an unstable state in which the memory can be modified and reconsolidated (Dudai, 2006; Elsey et al., 2018; Lee et al., 2017; Nader & Hardt, 2009). The cycle of reactivation and reconsolidation could occur more than once and constitutes a crucial part of the learning flexibility in daily experience. Interestingly, recent studies have shown that reactivation could facilitate the perceptual learning of texture discrimination task (Amar-Halpert et al., 2017; Chen & de Beeck, 2021; Klorfeld-Auslender et al., 2022). The strengthened memory through the reactivation and reconsolidation cycles was proposed to account for the facilitation effect (Amar-Halpert et al., 2017). Furthermore, the procedures based on reactivation were shown to resolve the interference of multiple learnings in motor learning (Herszage & Censor, 2017) and reward learning (Huang & Li, 2022b). These results were accompanied by the suggestion that memory integration triggered by the reactivated memory traces of the original learning and the novel memory of the new learning during the reconsolidation contributed to the prevention of the interference. This notion was consistent with the proposal that reactivation-induced integration of overlapping memory traces could facilitate the generalization of learning effect (Morton et al., 2017; Ritvo et al., 2019; Schlichting & Frankland, 2017). However, given the unique phenomenon of the location specificity in perceptual learning, it was unclear whether and how the reactivation-based procedure could facilitate the efficiency of multiple perceptual trainings by preventing interference between them. More specifically, how the location representation is involved in the integration process remained a critical issue to address.

The present study consisted of five psychophysical experiments to examine the role of memory reactivation and its induced memory integration in solving the potential interference between two perceptual learnings (Figure 1A). Experiments 1 established a baseline learning effect for the visual orientation discrimination task after five days of training (i.e., the first training). Experiment 2 demonstrated a proactive effect when a second training session at the untrained location was added two days after the first training. Experiment 3 showed that the interference could be eliminated by introducing a brief reactivation of the first training immediately before the second training. Experiment 4, in which a six-hour interval was introduced between the reactivation and the second training, showed that conducting the second training inside the reconsolidation window of the first training was critical for preventing the interference. Finally, in Experiment 5, we introduced a tilt discrimination task (TDT) after the reactivation but before the six-hour interval. This manipulation could also restore the effect of the second training but in a smaller magnitude, leading us to propose a two-level mechanism for the reactivation-induced memory integration in perceptual learning. We totally recruited 80 participants with 16 participants for each experiment. This is a larger sample size than typical multi-day perceptual learning studies to ensure sufficient statistical power when conducting between-subjects comparison. In a recent study (Hung & Carrasco, 2021) that examined the location specificity and transfer of the learning effect of the orientation discrimination task, nine participants were recruited for each training group and robust results were observed. Therefore, we believe that the sample size of the present study was appropriate to detect the potential training effects.

**Figure 1.**
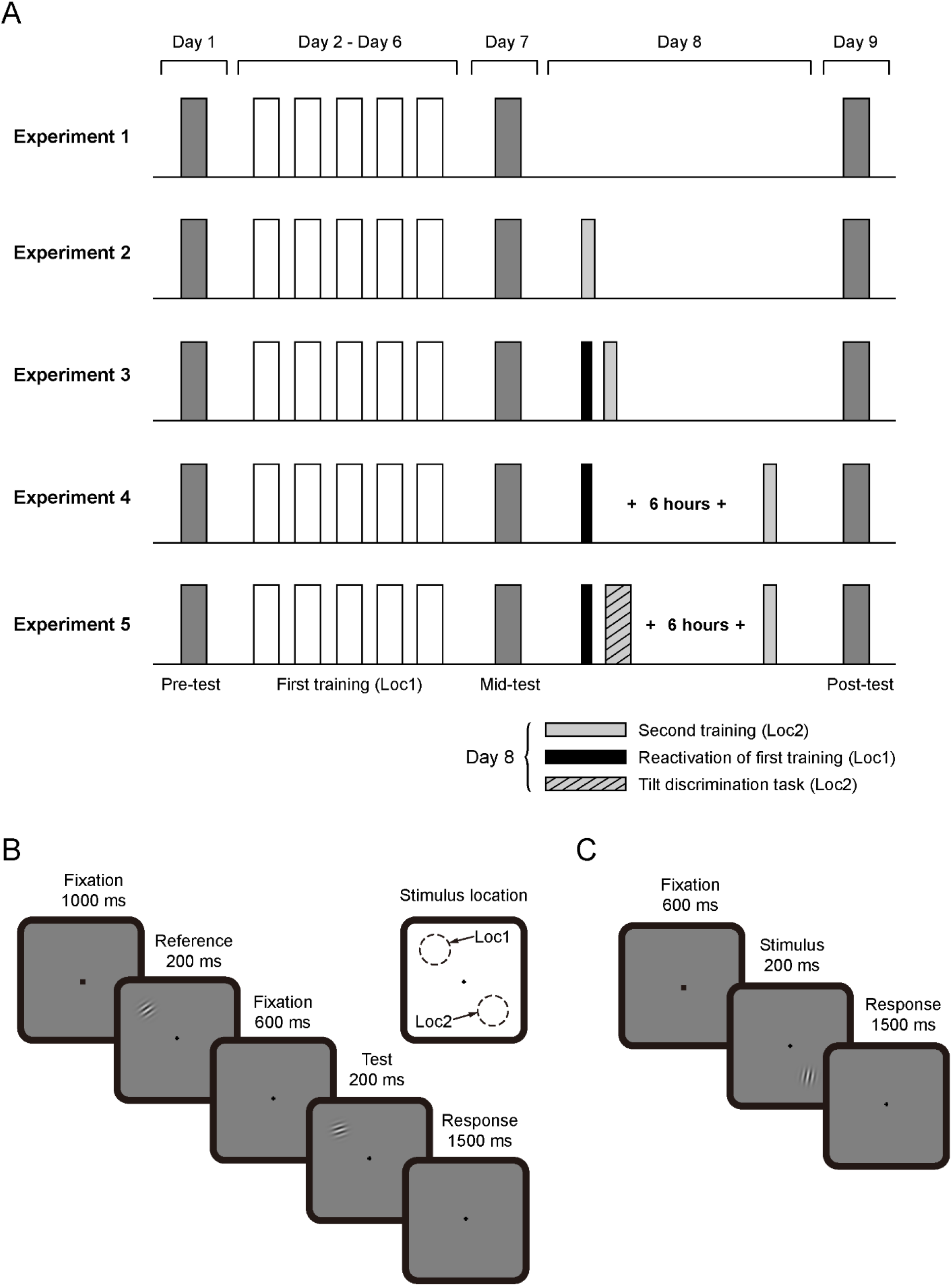
Experimental design. (A) Procedures of the experiments. (B) Orientation discrimination task. (C) Tilt discrimination task.

## Methods

### Participants

We totally recruited 80 participants with 16 participants from Peking University for each experiment (Experiment 1: 12 females, mean age = 19.50 years; Experiment 2: 11 females, mean age = 19.50 years; Experiment 3: 7 females, mean age = 19.69 years; Experiment 4: 11 females, mean age = 19.13 years; Experiment 5: 12 females, mean age = 19.75 years). Participants were with normal or corrected-to-normal vision and had no known neurological or visual disorders and provided written informed consent prior to the experiment. The local ethics committee approved the study.

### Stimuli

Stimuli were generated in MATLAB (MathWorks, Natick, MA, USA) using Psychtoolbox 3 package (Brainard, 1997; Pelli, 1997) and presented on a dark-gray background (~36 cd/m2) on a 20-inch CRT monitor (resolution of 1024 × 768 pixels and 60 Hz refresh rate). Gamma correction was applied to the monitor. Gabor stimuli (spatial frequency = 1.5 cycle/degree, contrast = 0.8, Gaussian filter sigma = 0.6, diameter ≈ 3.5°) were positioned in either upper-left or lower-right visual field with an eccentricity of 6.5°. The experiment was conducted in a dimly lit room. A chinrest was used to stabilize the head and maintain an 85 cm viewing distance.

### Procedure

The present study consisted of five psychophysical experiments (Figure 1A). All experiments were completed within nine days that consisted of a pre-test session (day 1), first training sessions (days 2-6), a mid-test session (day 7), and a post-test session (day 9). The experiments differed from each other in day 8.

#### Orientation discrimination task

As shown in Figure 1B, each trial of the orientation discrimination task began with a 1000 ms fixation interval, followed by the presentation of the reference Gabor stimulus for 200 ms at periphery (upper-left or lower-right). After another fixation interval of 600 ms, the test Gabor stimulus was shown for 200 ms at the same location as the reference. The participants were instructed to judge whether the test stimulus was tilted clockwise or counter-clockwise relative to the reference stimulus with a keyboard within 1500 ms from the offset of the test stimulus. The base orientation of the reference-test pair was 55° or 145°, counterbalanced across participants. One stimulus in the pair was randomly assigned to the base orientation and the other stimulus’ orientation was the base orientation plus a deviation. The orientation difference (i.e., the deviation) between reference and test stimuli was controlled by a three-down one-up staircase method (step size = 0.5°). This method converged to 79.4% correct responses. For the first block of each condition, the initial orientation difference was 5°. The stopping criterion of a block was 15 reversals or 100 trials, whichever was met first. The threshold in each block was determined by the mean orientation difference of the last eight reversals. The initial orientation difference of the subsequent blocks was set to be the threshold of the last block of the same condition.

#### Test sessions

In each test session, participants completed three runs of the orientation discrimination task. Each run consisted four stimulus conditions with one block for each condition. The four conditions were Loc1–Ori1, Loc1–Ori2, Loc2–Ori1, and Loc2–Ori2 (Loc1 was the location for the first training, Loc2 was the location for the second training, Ori1 was the stimulus orientation for both trainings). The order of the four conditions in each run was randomized. There was no feedback during the test sessions. There was a practice of 120 trials before the pre-test session, with 30 trials for each condition. Auditory feedback was provided after incorrect choices in the practice.

#### Training sessions

Participants were trained on the orientation discrimination task with Gabors presented at the same orientation and location throughout training. The stimulus conditions for the first and second trainings were Loc1–Ori1 and Loc2–Ori1, respectively. The orientation (55° or 145°) and location (upper-left or lower-right) of the trained Gabor stimulus were counterbalanced across participants. For the first training at Loc1, each participant was trained for five days with one session in each day. Each session consisted of 16 blocks. For the second training of day 8 at Loc2, each participant was trained for 8 blocks. There was auditory feedback after incorrect choices. The threshold of each training session was determined by the mean of the thresholds from the session.

#### Reactivation session

The reactivation session was identical to the first training session except that the participants were only trained for 2 blocks.

#### Tilt discrimination task (TDT)

As shown in Figure 1C, each TDT trial began with a 1000 ms fixation interval, followed by the presentation of a Gabor stimulus at Loc2 for 200 ms. The participants were required to judge whether the test stimulus was tilted clockwise or counter-clockwise relative to the vertical with a keyboard within 1500 ms from the offset of the test stimulus. There were five deviations from the vertical: −2.25°, −1.5°, −0.75°, 0°, 0.75°, 1.5°, and 2.25°. There was a practice of 28 trials at the beginning of the TDT session. The formal experiment of the session consisted of 4 blocks, with 140 trials in each block.

### Data analysis

The present study focuses on the two trained stimulus conditions (Loc1–Ori1 and Loc2–Ori1) and the influence of reactivation manipulations on the training effects of the two conditions. Therefore, the main results are shown only for these two conditions of interest (referred as Loc1 and Loc2 because they shared the same orientation) to avoid the complexity of conducting too many comparisons in statistics.

Repeated measures two-way ANOVA on threshold, with session (pre-test and mid-test) and location (Loc1 and Loc2) as factors, was performed to examine the effect of the first training. Repeated measures two-way ANOVA on threshold, with session (mid-test and post-test) and location (Loc1 and Loc2) as factors, was performed to examine the effect of the second training.

To reduce the impact of initial threshold on the training effect in between-experiment comparison, we calculated mean percent improvement (MPI) to evaluate the performance improvements between test sessions. MPI_pre-mid_ was defined as ((pre-test threshold – mid-test threshold)/pre-test threshold) × 100 to measure the improvement from pre-test to mid-test session. MPI_mid-post_ was defined in a similar fashion for the improvement from mid-test to post-test session. Mixed ANOVAs with experiment as a between-subjects factor and location (Loc1 and Loc2) as a within-subjects factor were performed to compared the training effect across experiments.

## Results

### Experiment 1

Experiment 1 served as a baseline for the training effect of the procedure (Figure 2). Training improved participants’ discrimination performance, as revealed by the decreased threshold from the first session (mean = 2.36°, SD = 0.629°) to the fifth session (mean = 1.96°, SD = 0.477°) of the training phase (paired *t* test: *t*(15) = 4.36, *p* = 0.001, Cohen’s *d* = 1.09). The training effect was further evidenced by the significant main effect of session (*F*(1,15) = 53.14, *p* < 0.001, 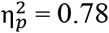) when we compared the estimated discrimination thresholds at the trained location (Loc1) and an untrained location reserved for the second training (Loc2) before (pre-test) and after (mid-test) the training sessions. The training effect was largely specific to Loc1 as revealed by the significant interaction between session and location (*F*(1,15) = 6.17, *p* = 0.025, 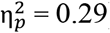). The thresholds in the two locations were statistically indistinguishable in the pre-test session (*t*(15) = 0.48, *p* = 0.64, Cohen’s *d* = 0.12). In mid-test session, the threshold at Loc1 was significantly lower than that of Loc2 (*t*(15) = 2.97, *p* = 0.01, Cohen’s *d* =0.74).

**Figure 2.**
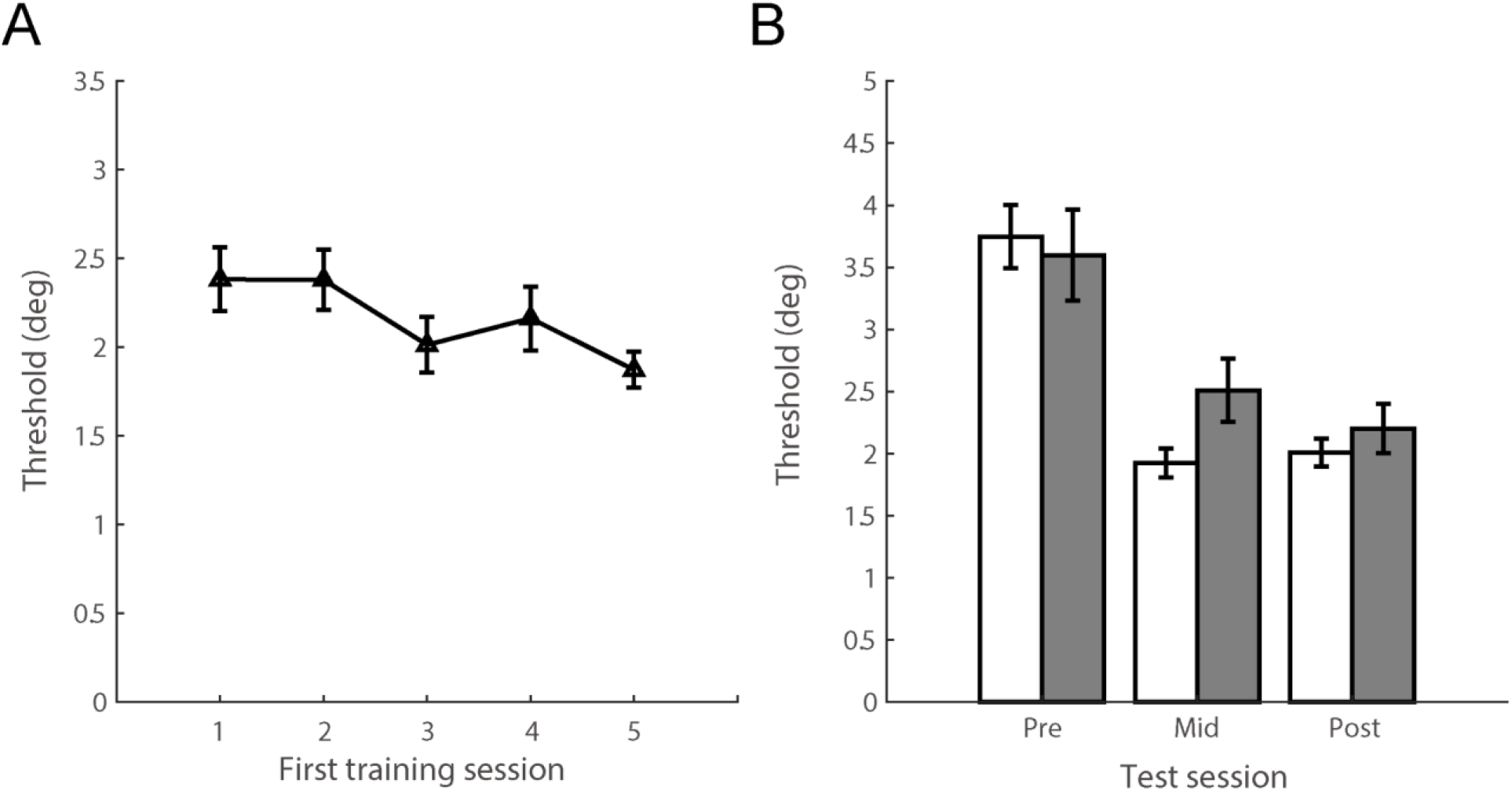
Experiment 1 results for (A) the first training sessions and (B) the test sessions. Error bars represent standard errors.

### Experiment 2

Experiment 2 introduced a half session training of the same orientation at Loc2 and the results are shown in Figure 3. The second training was done two days after the first training to ensure that the first training was well-consolidated. The results showed no significant difference from Experiment 1 in the effects of the first training (MPI_pre-mid_, *F*(1,30) = 0.44, *p* = 0.514, 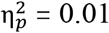). However, the second training at Loc2 was not effective as neither the threshold difference between the mid-test and post-test sessions (*F*(1,15) = 2.41, *p* = 0.141, 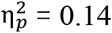) nor the interaction between session and location (*F*(1,15) = 1.68, *p* = 0.215 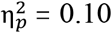) was significant. Also, the threshold changes from the mid-test to post-test sessions were not significantly different between the two experiments (*F*(1,30) = 0.01, *p* = 0.928, 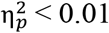). These results revealed a proactive interference effect from the first training to the second training, possibly due to the ten-fold difference in the amount of the training between them.

**Figure 3.**
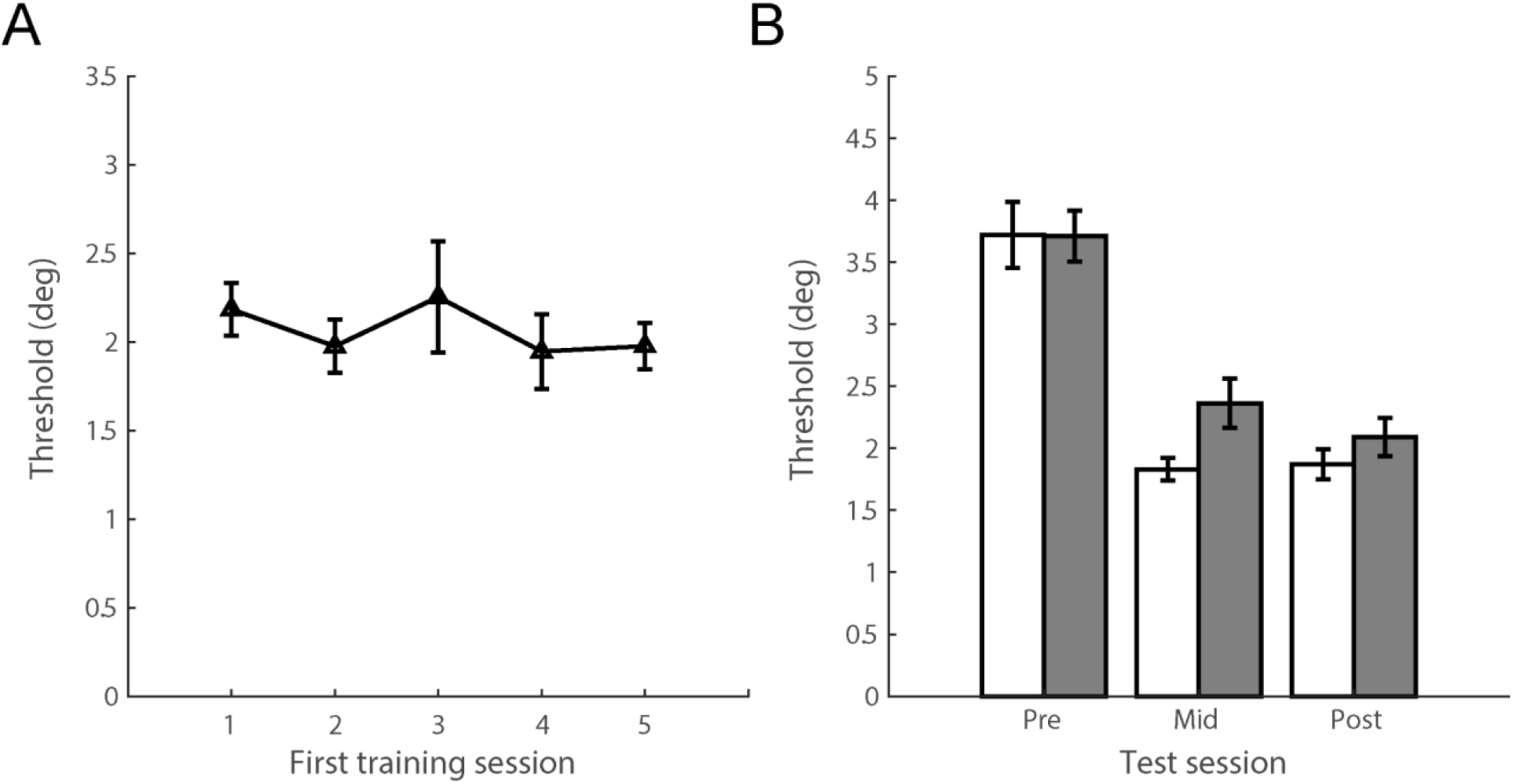
Experiment 2 results for (A) the first training sessions and (B) the test sessions. Error bars represent standard errors.

### Experiment 3

Huang and Li (2022) showed that conducting a new color-reward learning during the reconsolidation window of the reactivated old color-reward learning facilitated their integration and prevented them from interfering each other. This result agreed with the proposal that high degree of coactivation of two memory traces could lead to their integration and prevent the interference between them (Wammes et al., 2022). Therefore, we conducted Experiment 3 in which a brief reactivation of the first training was added before the second training. This manipulation would potentially induce the coactivation of the two trainings because second training would be temporally overlapped with the reconsolidation window of the first training. Indeed, as shown in Figure 4, the manipulation resulted in a significant effect for the second training (*F*(1,15) = 25.26, *p* < 0.001, 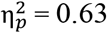), and this effect was largely specific to Loc2 (significant interaction between session and location: *F*(1,15) = 14.17, *p* = 0.002, 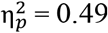). The threshold at Loc1 was statistically lower than that of Loc2 in the mid-test session (*t*(15) = 3.75, *p* = 0.002, Cohen’s *d* = 0.94). However, in post-test session, the thresholds at the two locations were statistically indistinguishable (*t*(15) = 0.82, *p* = 0.426, Cohen’s *d* = 0.21). Further between-experiments comparisons in MPI_mid-post_ revealed that the effects of the second training were significantly larger in Experiment 3 as compared with either Experiment 1 (*F*(1,30) = 14.31, *p* = 0.001, 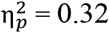) or Experiment 2 (*F*(1,30) = 18.84, *p* < 0.001, 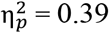), suggesting that combining the reactivation with the second training successfully release the inhibition of learning at the previously untrained location (i.e., Loc2). Additionally, the performance at Loc1 was further improved after the second training (one-sample t-test against 0 for the MPI_mid-post_: *t*(15) = 2.76, *p* = 0.015, Cohen’s *d* = 0.69).

**Figure 4.**
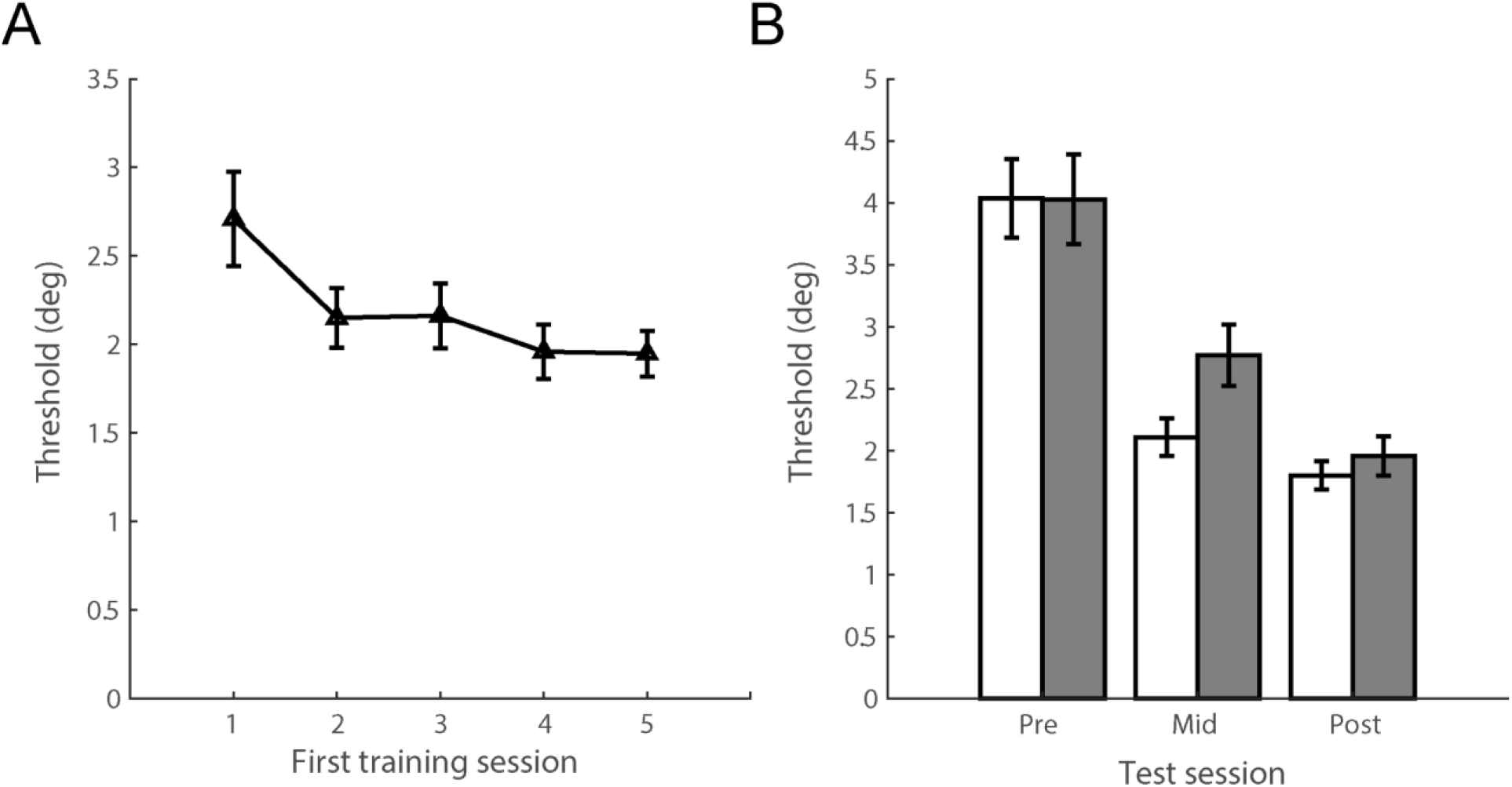
Experiment 3 results for (A) the first training sessions and (B) the test sessions. Error bars represent standard errors.

### Experiment 4

To validate that the coactivation of the two trainings was the key factor in restoring the effect of the second training, we conducted Experiment 4 in which a six-hour interval was placed between the reactivation and second training sessions (Figure 5). We chose six hours as the time window for the process of reconsolidation based on previous literatures (Bang et al., 2018; Huang et al., 2022; Huang & Li, 2022b; Nader et al., 2000). Because the second training was conducted after the first training’s reconsolidation, no coactivation of the two trainings would occur and their integration was not possible. The results revealed no significant effect of the second training (*F*(1,15) = 1.15, *p* = 0.301, 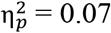). The effects of the second training as defined by MPI_mid-post_ were not significantly different from those of Experiment 2 (*F*(1,30) = 1.47, *p* = 0.235, 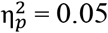), but were significantly smaller than those of Experiment 3 (*F*(1,30) = 16.34, *p* < 0.001, 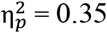). Re-emergence of the proactive interference after introducing this six-hour interval confirmed the important role of reactivation-induced memory integration in Experiment 3.

**Figure 5.**
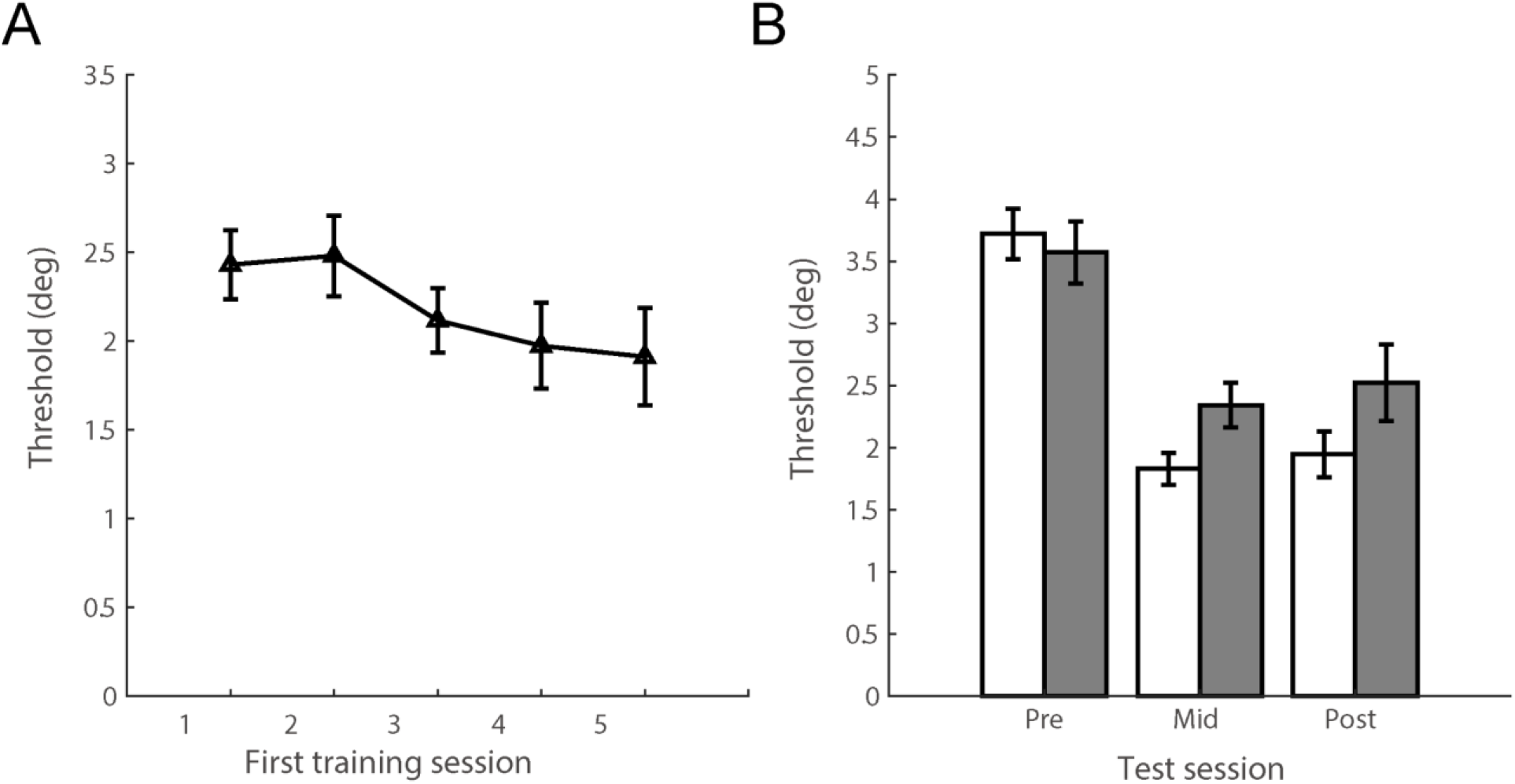
Experiment 4 results for (A) the first training sessions and (B) the test sessions. Error bars represent standard errors.

### Experiment 5

We then asked what was the role of location representation in the observed proactive interference in Experiment 2. Simply paying attention to Loc2 seemed not sufficient to solve the interference because the second training of Experiment 2 directed the participants’ attention to Loc2 and yet no learning occurred. It was the learning at Loc2 that was inhibited by the first training, potentially due to the learned low priority of attention during the first training (Huang & Li, 2022a; Wang & Theeuwes, 2018). Experiment 3 demonstrated an effective approach to foster the integration of the two trainings and thus release the inhibition of new learning caused by the intensive first training. Because the two trainings differed only at their stimulus locations, we would expect that the coactivation of the representations of the two locations was a part of the integration process. To test this hypothesis, we conducted Experiment 5 in which a tilt discrimination task at Loc2 was performed immediately after the reactivation and before the six-hour interval. With this setting, any difference in performance change following the second training between Experiments 4 and 5 would be attributed to the addition of the tilt discrimination task. We expected that adding the tilt discrimination task would integrate the two locations and release the inhibition of new learning at Loc2. Indeed, we found that the effects of the second training were restored (*F*(1,15) = 17.87, *p* = 0.001, 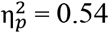; Figure 6), as the training effects defined by MPI_mid-post_ were significantly larger than those of Experiment 4 (*F*(1,30) = 4.81, *p* = 0.036, 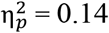), suggesting that the location-based learned attentional suppression may play an important role in the observed proactive interference. However, the restored effects were significantly smaller than those of Experiment 3 (*F*(1,30) = 6.95, *p* = 0.013, 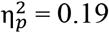) and there was no further improvement at Loc1 (one-sample t-test against 0 for the MPI_mid-post_: *t*(15) = 0.10, *p* = 0.919, Cohen’s *d* = 0.03) after the second training, suggesting that only location representations were integrated to unlock the inhibition of learning at Loc2 in Experiment 5.

**Figure 6.**
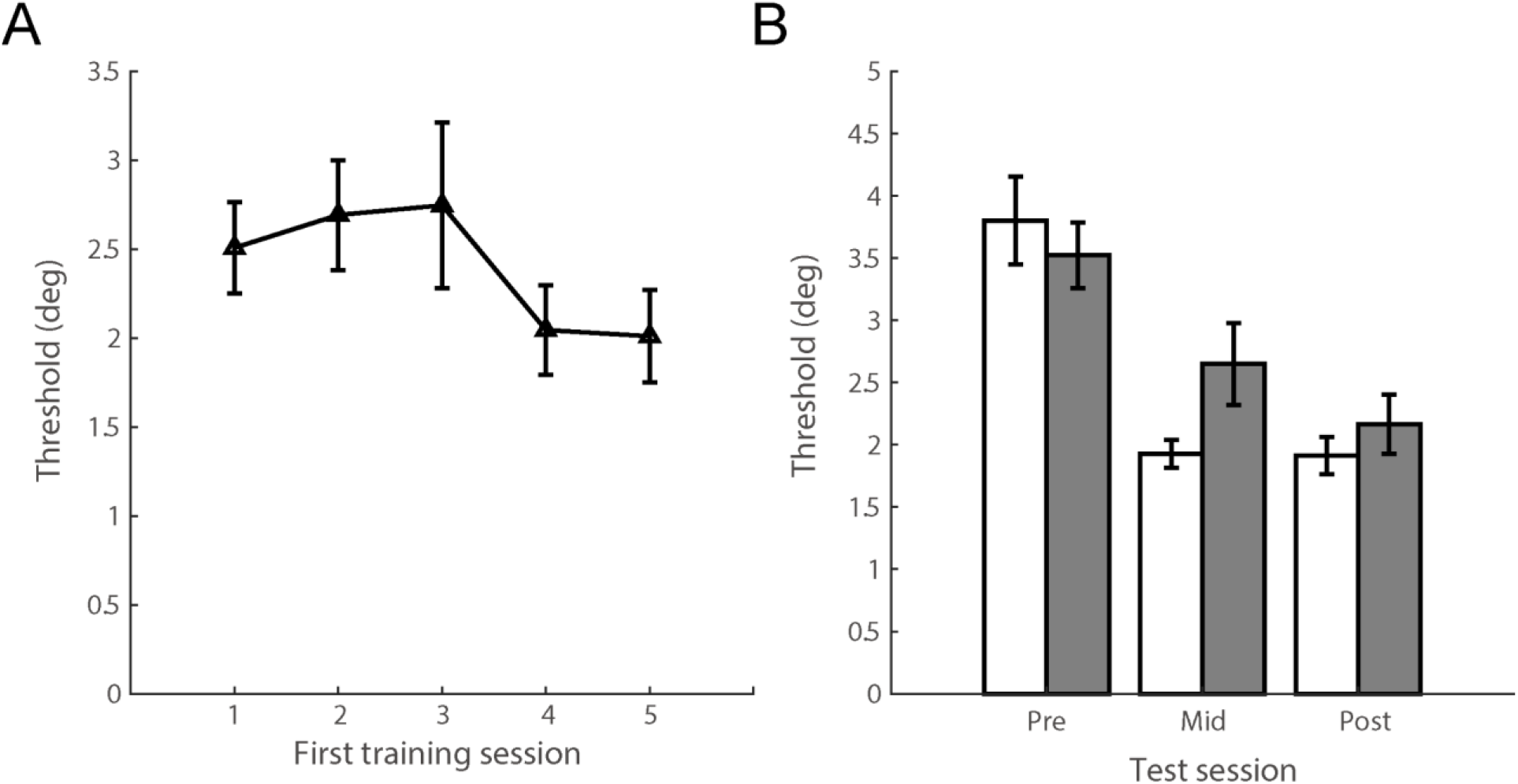
Experiment 5 results for (A) the first training sessions and (B) the test sessions. Error bars represent standard errors.

## Discussion

Summary of the results across experiments is shown in Figure 7. The reactivation manipulations in Experiments 3 and 5 revealed two related but different results (Figure 7B). In Experiment 3, the reactivation of the first training was immediately before the second training, resulting in significant training effect at Loc2 and additional improvement at Loc1. In Experiment 5, the reactivation of the first training was immediately followed by a tilt discrimination task at Loc2 and the second training was conducted six hours later. The second training could also be protected from the proactive interference in this procedure but the additional performance improvement at Loc1 was not observed. Importantly, the performance improvement from the mid-test to post-test session was smaller in Experiment 5 as compared with that of Experiment 3. We suggest a two-level mechanism of reactivation-induced memory integration to account for these results.

**Figure 7.**
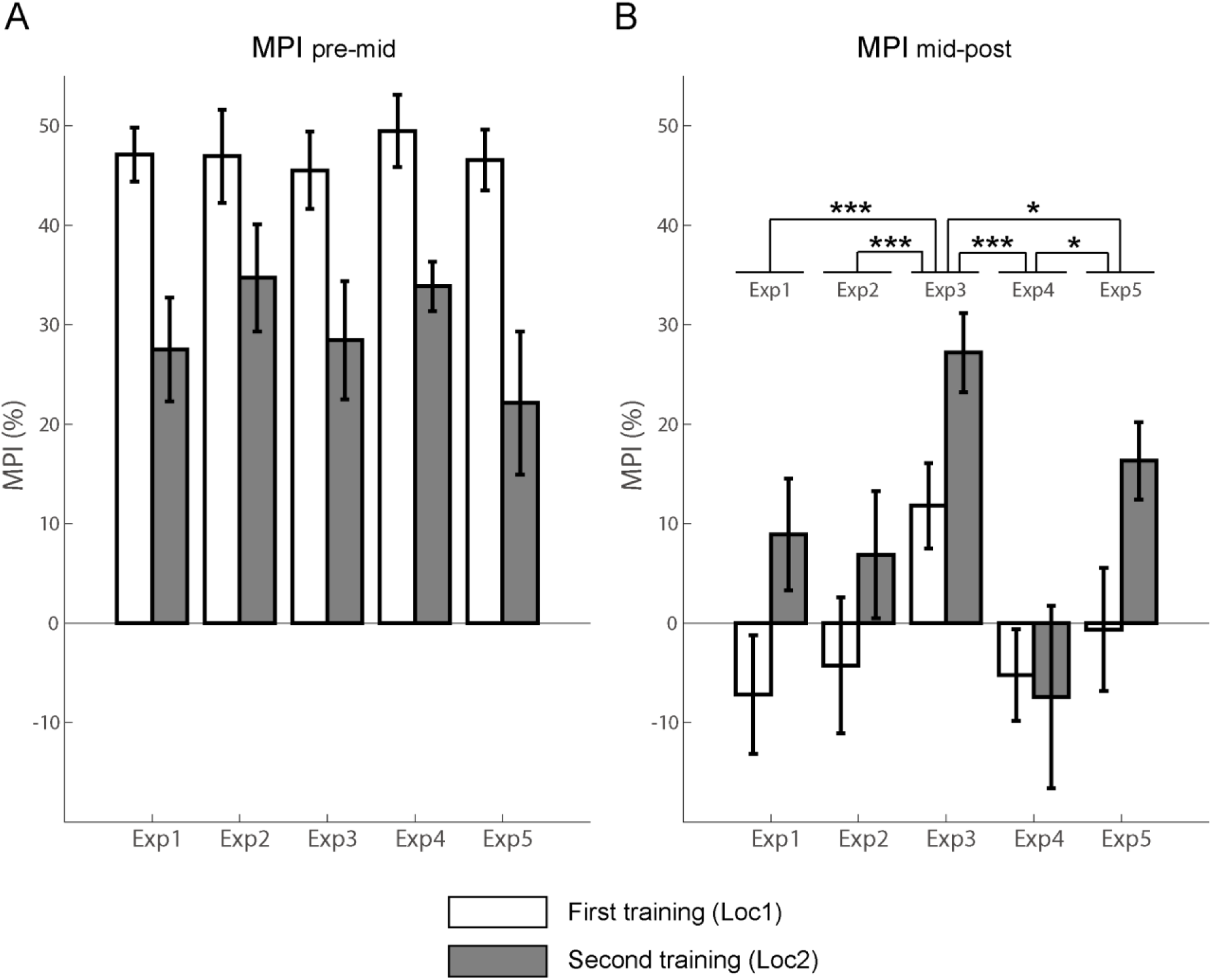
Summary of the results. Mean percent improvements (MPIs) after the (A) first training (MPI_pre-mid_) and (B) second training (MPI_mid-post_) are shown for each experiment. Error bars represent standard errors. * *p* < 0.05, *** *p* < 0.001.

The first level of the mechanism is to release the inhibition of new learning at the previously untrained location (Loc2) by integrating the two locations during the reconsolidation process. This can be achieved by the coactivation of the two locations when the location representation of Loc2 was activated during the reconsolidation window of the reactivated first training at Loc1. In the present study, the representation of Loc2 was activated by the second training and the tilt discrimination task in Experiment 3 and Experiment 5, respectively. Accordingly, we have observed significant improvement at Loc2 following the second training in both experiments. However, only releasing the inhibition of new learning at Loc2 in Experiment 5 was not sufficient to further elevate the performance at Loc1. We would attribute the additional improvement at Loc1 following the second training in Experiment 3 to the second level of the mechanism. In the second level, the memory traces of the trainings at the two locations were integrated due to their coactivation that occurred when the second training was conducted during the reconsolidation of reactivated first training. This process could form an integrated memory of the two trainings so that training at Loc2 would benefit the performance at both locations as was evident in Experiment 3. Together, these results suggest that which level of the mechanism dominates the reactivation-induced integration process depends on the representations that are coactivated during the reconsolidation.

A recent theory named nonmonotonic plasticity hypothesis (NMPH) suggests that coactivation of two overlapping memories could lead to memory integration or differentiation depending on the level of the coactivation (Ritvo et al., 2019; Wammes et al., 2022). NMPH states that the two memories would be integrated if they are strongly coactivated. Otherwise, if one memory is strongly activated and the unique components of the other memory are moderately active, the coactivation would be moderate one and result in differentiation of the two memories. In Experiment 3, to reactivate the first training, we simply had the participants performing the training task that differed from the second training only at the stimulus location. Hence, the activation of the first training during its reconsolidation was most likely at a high level given the concurrently performed second training. Similarly, the representations of the two locations were also likely to be strongly coactivated in Experiment 5. In both experiments, NMPH and other models of memory integration (Morton et al., 2017; Schlichting & Frankland, 2017) would predict the occurrence of integration that agreed with the observed results.

The first training showed partial transfer of learning effect to the untrained location in all experiments (*ps* < 0.01; Figure 7A). This was in sharp contrast with the blocking of the further learning at the untrained location in Experiment 2. Previous studies have shown that the transfer of learning to untrained locations can be elevated by double training method that was suggested to release the attentional suppression to the untrained location by performing an unrelated task at that location (Wang et al., 2014; Xiao et al., 2008; Zhang et al., 2010). In these studies, the unrelated task was generally performed for several sessions in order to release the learned attentional suppression. This setting was consistent with the results in Experiment 2 in which a short training session at the untrained location was not sufficient to release the inhibition of new learning. However, by adding a brief activation of the first training before the second training, we enabled the new learning at the untrained location and induced further improvement for the first training, demonstrating that reactivation-induced memory integration is a practical way to foster the efficiency of multiple perceptual learnings. Further, our results also validated the idea that reactivation-induced integration could facilitate the generalization of learning (Morton et al., 2017; Ritvo et al., 2019; Schlichting & Frankland, 2017).

Our results suggest an encoding phase hypothesis that the integration occurred during the second training and does not support a retrieval hypothesis that pattern separation was induced by the reactivation so to prevent interference at the retrieval stage. If pattern separation was realized during the reconsolidation process, it would likely to generate two separate associations for the two locations, and thus prevent their interference during the retrieval (Huang et al., 2022). However, the retrieval hypothesis cannot explain the additional performance improvement at Loc1 after the reactivation and second training in Experiment 3. If the two trainings were dissociated during the reconsolidation, the second training would not benefit the performance at the location of the first training. Considering that contextual cueing could facilitate pattern separation and prevent interference of perceptual learning (Huang et al., 2022), we suggest that both the pattern separation (memory differentiation) and memory integration could benefit the efficiency of multiple perceptual learnings. The difference was the ways that memory differentiation and memory integration were induced, as contextual cueing could promote the former and reactivation plus new learning promotes the latter.

In summary, the present results suggest a two-level mechanism for reactivation-induced memory integration and its roles in preventing proactive interference in perceptual learning. These findings could be adopted in future applications of various forms of learning that suffer from the interference from previous learning experience.

## Notes

Author note: The authors declare no competing financial interests. This work was supported by grants from Science and Technology Innovation 2030 - Brain Science and Brain-Inspired Intelligence Project (2021ZD0200204).

### Competing Interest Statement

The authors have declared no competing interest.

### Summary of Updates

Figure 2 is split into 5 figures (Fig.2 to Fig.6), one for each experiment. Figure 3 is now Figure 7. Sections on abstract, introduction, and discussion are updated to extend the theoretical implication of the study. Section on methods is updated to clarify a few parameters of the stimuli.

